# Targeted deletion of *Nmnat1* in mouse retina leads to early severe retinal dystrophy

**DOI:** 10.1101/210757

**Authors:** Xiaolin Wang, Yu Fang, Rongsheng Liao, Tao Wang

## Abstract

Mutations in *NMNAT1* can lead to a very severe type of retinal dystrophy, Leber congenital amaurosis, in human patients, characterized by infantile-onset or congenital retinal dystrophy and childhood blindness. The loss-of-function mouse models of *Nmnat1* have not been well-established, since the complete knock-out (KO) of *Nmnat1* in mice results in embryonic lethality. Here, we generated retina-specific KO by using the Crxpromotor-driving Cre combined with the flox allele. By a panel of histological and functional analyses, we found that *Nmnat1* conditional KO (cKO) mice have early severe retinal dystrophy. Specifically, the photoreceptors of *Nmnat1* cKO mice are almost diminished and the retinal functions also become completely abolished. Our results established a loss-of-function model for *Nmnat1* in mice, which will be useful for studying the detailed functions of NMNAT1 in the retina.

## Introduction

Leber congenital amaurosis (LCA) is a severe inherited eye disease characterized by infantile-onset visual impairment and vision loss (1, 2). Though LCA is a rare disease with an incidence about only 1 in 80000 (3), it is the most common cause of incurable children blindness (10%-18%) (1). LCA is a highly genetically heterogeneous disease. During the past 20 years, various genetic analyses have identified at least 19 genes that cause LCA(4–27). However, the mutations in these genes account for approximately 70% of all LCA cases, leaving 20%-30% unsolved cases to be discovered for their genetic basis (28).

Sequencing studies have recently found that mutations in *NMNAT1* can cause LCA and these constitute approximately 10% of unsolved cases (28–47). The majority of *NMNAT1* mutations found in LCA patients are missense, with a small portion are nonsense and frameshift. Most of the patients share a variant (c.769G>A, E257K), while the other allele is heterogeneous spreading from N-terminal to C-terminal (30). *In silico* analysis predicts that these mutations will affect the enzymatic activity, hexamerization, hydrophobic interactions or nuclear localization of NMNAT1 protein (31).

Though one mouse model with *Nmnat1* homozygous missense mutation was reported recently (48), it remains unknown what is the impact of retina-specific deletion of *Nmnat1* in mice. Therefore, in this study, we generated retina-specific knock-out *Nmnat1* mice and found that this genetic defect can lead to early severe retinal dystrophy.

## Methods

### Mouse study

*Nmnat1* retina-specific KO mice were generated by crossing the *Nmnat1* ^flox/flox^ mice with Crx-Cre ^+/−^ mice to get the *Nmnat1* ^flox/+^; Crx-Cre ^+/−^mice. Then these mice underwent further mating to get the *Nmnat1* retina-specific KO mice. The mouse study protocols were approved by the Institutional Review Board of Sichuan Medical College.

### Histological study

For histological studies of the mouse retina, the mice were sacrificed and the eyecups were enucleated and fixed. The eyecups were embedded in paraffin and then sectioned to 10om slices. The sections underwent Hematoxylin and Eosin (HE) staining after rehydration. The number of nuclear layers were counted as the indicator of retinal thickness.

### Electroretinogram (ERG)

As for the ERG experiments, the mice underwent dark adaptation for overnight (at least 10 hours) before the experiments, and the ERGs are recorded in the dark with dim red lights. Mice were anesthetized with Ketamine and then put in the chambers for the actual testing. The heads of the mice were stabilized properly. Six different light stimuli amplitudes were applied and the retinal responses were recorded by the electrodes attached to the cornea of the mice. The responses contain a-wave and b-wave responses. Then the data were analyzed and plotted according to their genotypes.

## Results

At the age of P7, since the outer nuclear layer (ONL) and inner nuclear layer (INL) have not been fully developed and separated, we used the thickness of the two layers as the indicator. By H&E staining, we found that the ONL and INL thickness in those cKO mice is only about 45% compared with that in WT mice. The thickness of ganglion cell layer (GCL) is a little bit smaller in cKO mice compared with WT (Figure 1A).

**Figure 1.**
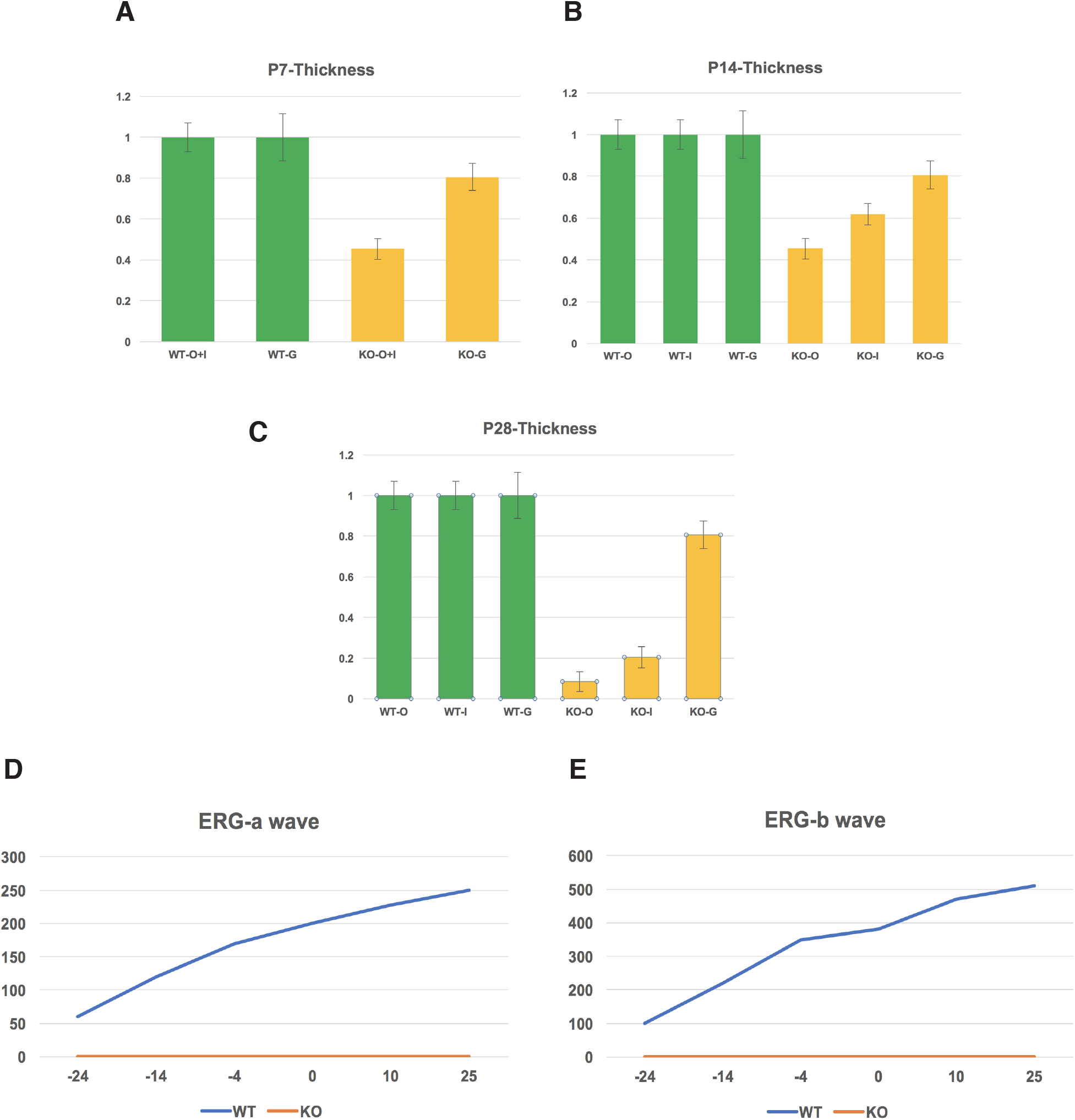
(A) The relative retinal thickness of retinal cell layers at P_7_ in WT and KO mice (n=5) (B) The relative retinal thickness of retinal cell layers at P_14_ in WT and KO mice (n=6). (C) The relative retinal thickness of retinal cell layers at P_28_ in WT and KO mice (n=6). (D) The ERG a-wave responses at P_28_ in WT and KO mice. (E) The ERG b-wave responses at P_28_ in WT and KO mice. O+I: outer nuclear layer and inner nuclear layer; G, ganglion cell layer. The unit in ERG experiments is μV.

When the mice grow to P14, the retinal layer thinning becomes more remarkable in cKO mice compared with WT mice. Both the ONL and INL show apparent dystrophy in cKO mice. Similar to P7, the GCL is about 20% thinner in cKO mice (Figure 1B).

At the age of P28, the ONL (photoreceptor layer) becomes almost diminished in cKO mice, and the INL is also severely dystrophic in cKO. The GCL does not show apparent dystrophy in cKO, similar to P7 and P14 (Figure 1C).

As for the functional analysis, we used ERG to measure the retinal electric responses to the light stimuli. At the age of P28, ERG experiments revealed that the cKO mice do not have any a-wave and b-wave responses to light compared with WT mice. (Figure 1D and 1E). These results demonstrated the complete abolishment of retinal functions in cKO mice upon NMNAT1 deficiency.

## Discussion

Previous studies on axon protection indicated that NAD+ biosynthesis activity is essential for NMNAT1’s role in protecting axons from degeneration (49–57). NAD^+^ is a fundamental molecule for the living organisms. It is a coenzyme involved in various cellular processes. NAD/NADH redox pair is essential for electric transfer chain and is utilized for the maintenance of cellular redox state (58). The axon protection conferred by NMNAT1 seems to be closely related to mitochondria, which is a critical organelle for the maintenance of redox state (59, 60). In addition, NMNAT1 also has other functional implications(50, 59, 61–78).

Interestingly, most of the LCA patients with *NMNAT1* mutations show specific degeneration in the macula where the redox homeostasis is particularly important due to high oxygen consumption conferred by high density of photoreceptors (30, 79–83). This evidence, with the NAD+ function described early, provides a strong linkage between the need for NAD^+^ biosynthesis and the maintenance of photoreceptor normal function.

*Nmnat1* complete knockout mice have been generated but they are embryonically lethal (66). Therefore, for studying the function of NMNAT1 in retina, conditional KO mice specifically targeting photoreceptors (NMNAT1 flox/flox; Crx-Cre+/-) were generated by us. Our results show that *Nmnat1*-cKO mice show extremely rapid photoreceptor loss and blindness. The phenotypes are somewhat consistent with some human LCA patients who were blind at very early stages, thus presenting a useful model for NMNAT1 functional study in the retina in the future.

## Acknowledgements

This work was supported by Sichuan Province Nature Science Funding (No.31600642). We thank Dr. Fei Luo for the help on study design.

